# Novel *bla*_KPC_-carrying species identified in the hospital environment

**DOI:** 10.1101/069856

**Authors:** Hang T.T. Phan, Zoie Aiken, Oluwafemi Akinremi, Julie Cawthorne, Andrew Dodgson, Angela Downes, Matthew Ellington, Ryan George, Amy Mathers, Nicole Stoesser, Stephanie Thomas, on behalf of the TRACE Investigators’ Group

## Abstract

*bla*_KPC_, encoding one of five dominant global carbapenemase families, is increasingly identified in environmental species difficult to characterize using routine diagnostic methods, with epidemiological and clinical implications. During environmental hospital infection prevention and control investigations (Manchester, UK) we used whole genome sequencing to confirm species identification for isolates infrequently associated with *bla*_KPC_ and/or difficult to classify by MALDI-ToF. Four previously undescribed *bla*_KPC_-carrying species were identified from the hospital environment, including a putative, novel *Enterobacter* species.

## Main text

Carbapenems are used to treat serious infections resistant to other antimicrobial agents. *Klebsiella pneumoniae* carbapenemases (KPC), encoded by *bla*_KPC_ alleles are a major carbapenemase family, conferring resistance to most beta-lactams. *bla*_KPC_, first identified retrospectively in *K. pneumoniae* dating from 1996, has since been observed in other Enterobacteriaceae, Pseudomonadaceae and *Acinetobacter* spp.[1]. The host species range of *bla*_KPC_ has not been defined, and may be more diverse than originally thought, given its presence in a number of environmental reservoirs[2-4].

Accurate species identification is important for epidemiological and clinical reasons. Species identification methods (e.g. biochemical profiling, Matrix Assisted Laser Desorption Ionization Time-of-Flight [MALDI-ToF]), or partial sequence-based typing (e.g. 16S rRNA gene sequencing) can be inconsistent, particularly for Enterobacteriaceae[5, 6]. Whole genome sequencing (WGS) has an advantage over these methods in using the complete genetic content of an organism for identification[7].

Here, WGS was used to confirm species identification of KPC-producers cultured during environmental infection prevention and control investigations undertaken on wards with KPC-positive patients in two hospitals in Manchester, UK (May 2012-March 2013, April 2015). High-touch surfaces, toilet edges and sinks were sampled with EZ-Reach sponges squeezed out into neutralizing buffer; water samples were collected from sink drain P-traps with plastic tubing and a syringe. Samples were centrifuged, and pellets sub-cultured overnight in tryptic soy broth (5mls) with ertapenem 10μg discs. Broths were sub-cultured on chromID CARBA plates (bioMérieux, Marcy-l’Étoile, France); different Enterobacteriaceae colony morphotypes were confirmed as *bla*_KPC_ positive by PCR and species determined by MALDI-ToF (Bruker, Billerica, Massachusetts, United States).

WGS was performed on 142 isolates from 55 samples using the Illumina HiSeq platform (150bp, paired-end reads). *De novo* assemblies were created using SPAdes 3.6[8], and assessed for integrity using REAPR[9]. BLASTn-based analyses against assemblies confirmed the presence of *bla*_KPC_ in all 142 isolates.

Species identification was consistent by both MALDI-ToF and WGS for 78 *Klebsiella pneumoniae* isolates, 11 *Enterobacter cloacae*, five *Escherichia coli*, five *Citrobacter freundii* and four *Klebsiella oxytoca*. However, limited sequence matches with the NCBI nr reference database were observed for two isolates characterized as *Escherichia vulneris* and/or *Enterobacter cancerogenus* by MALDI-ToF (9885_1 [81% sequence similarity/58% coverage with the top match]; and 9885_2 [84%/74%]), and 36 isolates characterized as *Pantoea agglomerans* by MALDI-ToF (all closely genetically related; only 88%/85% coverage with the top match). In addition, one isolate (1613625) was characterized as *Pluralibacter gergoviae* by MALDI-ToF and had a good sequence match to the NCBI nr reference database; however, this species had only been associated with *bla*_KPC_ in two previous disease-causing (i.e. not environmental) isolates (in Brazil and the USA)[10, 11]. We therefore further investigated the species identification of isolates 9885_1, 9885_2, 9885_4 (representative of the putative *P. agglomerans* group) and 1613625 using the genomic data.

Querying the assemblies against the Silva 16S database[12] (Table 1) demonstrated that each isolate had several hits above the suggested threshold for 16S-based species identification (>99.5% similarity). However, in all cases the first and second hits were divergent. Species identification was therefore undertaken using genome-wide average nucleotide identity (ANI) and alignment fraction (AF) scores, where ANI and AF scores 96-96.5% and 0.6 respectively are considered species-specific thresholds[7, 13]. Sequence assemblies were first annotated with PROKKA[14] and coding sequences were fed into the ANICalculator[13]. ANI scores for isolate assemblies and top hit species found by 16S analysis (where available as reference genomes or assemblies in GenBank; n=34) were calculated. These identified isolate 9885_1 as *vulneris*, 9885_4 as *Pantoea anthophila*, 1613625 as *P. gergoviae*, and 9885_2 as a presumptive new *Enterobacter* species (Table 1).

**Table 1.** Top 16S and genome wide species identification results of study *bla*_KPC_-positive isolates

| Isolate id | First hit species<br>(% identity) | Second hit species<br>(% identity) | Average nucleotide identity (ANI) species classification |  |  | Final species<br>classification |
| --- | --- | --- | --- | --- | --- | --- |
|  |  |  | Best species match | ANI<br>(bidirectional) | Alignment<br>Fraction (AF)<br>(bidirectional) |  |
| 9885_1 | *Escherichia*<br>*vulneris* (99.74) | *Enterobacter*<br>*hormaechei* (99.73) | *Escherichia vulneris* | 96.38\|96.37 | 0.79\|0.89 | *Escherichia*<br>*vulneris* |
| 9885_2 | *Citrobacter*<br>*murlinae* (99.80) | *Enterobacter*<br>*asburiae* LI (99.42) | *Enterobacter sp MGH24* | 86.69\|86.68 | 0.74\|0.80 | Novel<br>*Enterobacter* sp. |
| 9885_4 | *Pantoea*<br>*anthophila* (99.87) | *Pantoea*<br>*agglomerans* (99.87) | *Pantoea anthophila* | 98.75\|98.75 | 0.74\|0.96 | *Pantoea*<br>*anthophila* |
| 1613625 | *Enterobacter* sp.<br>*ETH-2*(100.00) | *Pluralibacter*<br>*gergoviae* (99.81) | *Pluralibacter gergoviae* | 98.91\|98.91 | 0.90\|0.89 | *Pluralibacter*<br>*gergoviae* |

We compared the genomes of these four isolates with those of closely related species (n=34) using a core/pan-genome approach (ROARY)[15], and including only those sub-groups with a core genome >200 core genes. For this analysis, contigs of plasmid origin (high sequence similarity with plasmids in the NCBI nr database) were excluded. A maximum likelihood phylogeny from core genome alignments was reconstructed; the core genome phylogenies (Figure 1) concurred with the ANI results and highlighted inconsistencies in traditional taxonomic assignations that are likely to be revised as more bacterial genome sequences become available[16].

**Figure 1.**
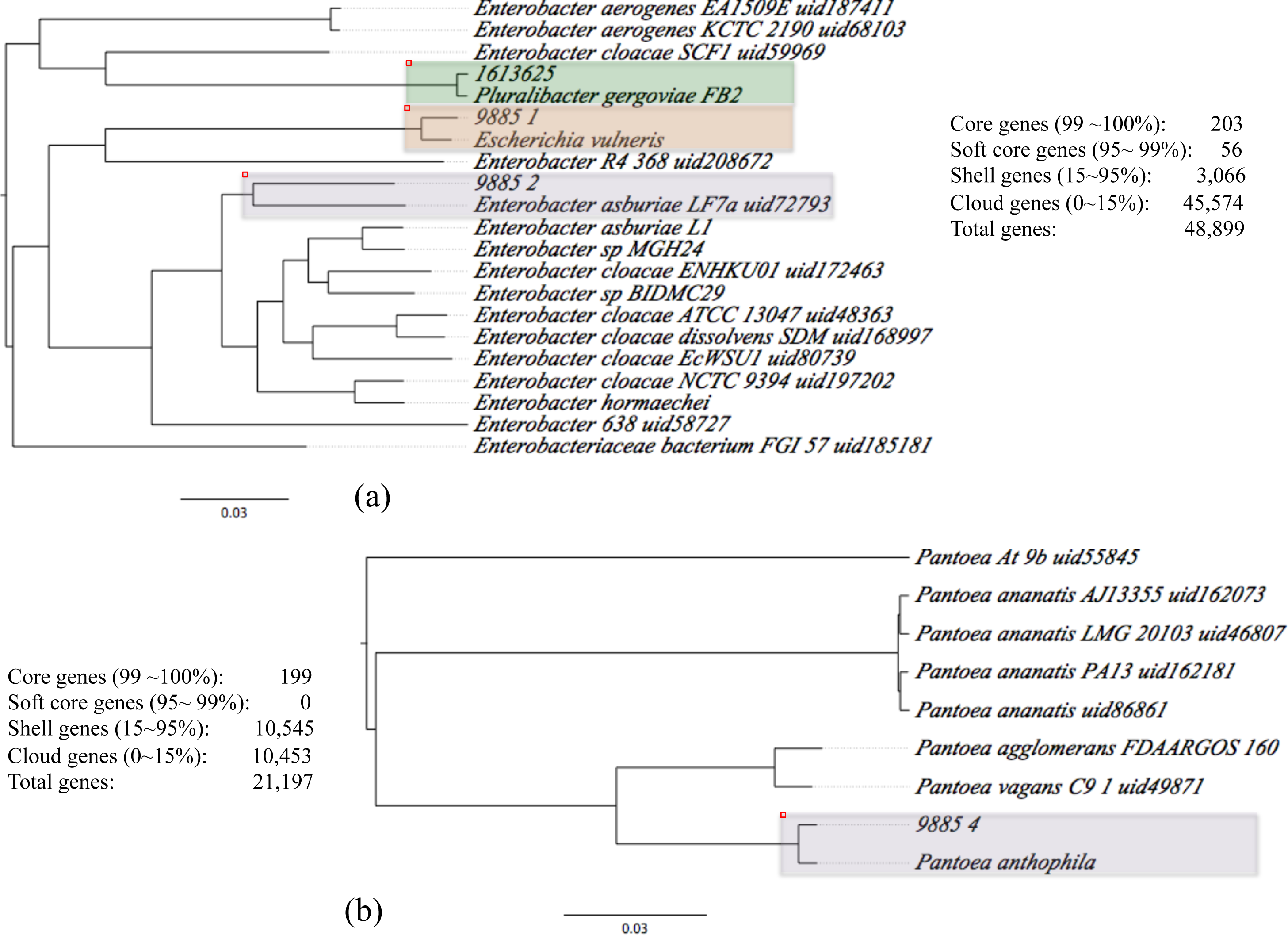
Core genome phylogenies of *bla*_KPC_-positive isolates investigated in detail in this study and of closely related reference genomes from NCBI/GenBank. (a) phylogeny of 9885_1, 9885_2 and 1613625 and reference *Enterobacter* spp. genomes; (b) phylogeny of 9885_4 and reference *Pantoea* spp. genomes

*E. vulneris* has been associated with both *Escherichia* and *Enterobacter* genera by DNA hybridization[17], perhaps explaining the initial identification of 9885_2 as either *E. vulneris/E. cancerogenus* by MALDI-ToF. *E. vulneris* has been described in wound infections, bacteraemia, peritonitis, urosepsis and meningitis[17-20], and its novel association with *bla*_KPC_ in this study is therefore potentially of clinical significance. *Pantoea* spp. are recognized as opportunistic pathogens and whilst *P. agglomerans* has been associated with *bla*_KPC_ previously[21], *P.anthophila*, found in lakes and diseased plants[22, 23], has not. Worryingly, in this survey, it was one of the most widespread *bla*_KPC_-positive environmental species (36 isolates from 8 sites from one ward, including clinical areas, staff areas and ward kitchens). *P. anthophila* may represent a successful intermediate species for the wider transmission of *bla*_KPC_ amongst more pathogenic strains within/between non-clinical/clinical sites. P. gergoviae has been isolated from a wide range of sources, including foodstuffs and insects[24, 25], but has not previously been identified with *bla*_KPC_ in environmental isolates. Of particular concern is the organism’s innate resistance to parabens, enabling it to exist as a contaminant in a variety of personal care products, including soaps and shampoos[25].

We have identified four species not previously associated with *bla*_KPC_ in the hospital environment, one a putative, novel *Enterobacter* sp. Accurate species identification of *bla*_KPC_-carrying environmental isolates has implications for understanding the wider reservoirs and transmission dynamics of this resistance gene family, assessing the pathogenic potential of these organisms, and for accurate susceptibility testing in isolates that have the potential to become clinically relevant. Although WGS is not currently widely used in routine diagnostics, the development of bench-top sequencers and falling costs may make this more feasible[26]. Species identification can also be hampered by incomplete reference genome databases, which is being addressed by the large-scale sequencing of type strain collections[27]. Our identification of *bla*_KPC_ in diverse species in the hospital environment supports previous data suggesting that this reservoir plays a role in the dissemination of *bla*_KPC_ across species and genera[2].

## Nucleotide sequence data

These have been deposited in GenBank under project number PRJNA324191, study number SRP076320 and accession numbers SRR3654271, SRR3654272, SRR3654273, SRR3654274.

## ACKNOWLEDGEMENTS

The Transmission of Carbapenemase-producing Enterobacteriaceae (TRAcE) study investigators are (alphabetical): Zoie Aiken, Oluwafemi Akinremi, Julie Cawthorne, Paul Cleary, Derrick Crook, Valerie Decraene, Andrew Dodgson, Matthew Ellington, Ryan George, Katie Hopkins, Rachel Jones, Cheryl Lenney, Amy Mathers, Ginny Moore, Sarah Neilson, Tim Peto, Hang Phan, Mark Regan, Anna C. Seale, Nicole Stoesser, Stephanie Thomas, Jay Turner-Gardner, Vicky Watts, Jimmy Walker, Sarah Walker, David Wyllie, William Welfare and Neil Woodford.

We are grateful to the Infection Control Teams and Microbiology Laboratory staff at the University Hospital of South Manchester NHS Foundation Trust and the Central Manchester University Hospitals NHS Foundation Trust; the Sequencing hub at the Wellcome Trust Center for Human Genetics, Oxford; and the Modernizing Medical Microbiology Informatics Group (MMMIG), Oxford.

